# Maternal den site fidelity of polar bears in western Hudson Bay

**DOI:** 10.1101/2024.01.09.574879

**Authors:** Natasha Klappstein, David McGeachy, Nicholas Pilfold, Nicholas Lunn, Andrew Derocher

## Abstract

Seasonal migrations allow to access temporally varying resources and individuals may show fidelity to specific locations. Polar bears (*Ursus maritimus*) are a sea ice dependent species that migrate between marine and terrestrial habitats, the latter being important for parturition and early cub rearing. However, fidelity to maternity den sites is poorly understood. We assessed polar bear maternal den site fidelity of the Western Hudson Bay subpopulation in Manitoba, Canada. Using capture and telemetry data collected between 1979 *−* 2018, we examined site fidelity from 188 maternity den locations from 78 individuals. We calculated within-individual inter-year distances between dens, and compared these to between-individual distances via non-parametric bootstrapping. We used generalised additive models to assess how maternal age, years between denning events, and sea ice conditions affected site fidelity. We found some evidence of site fidelity, as within-individual inter-year distances were smaller than between-individual den distances by approximately 18.5 km. As time between captures increased, inter-den distances also increased (ranging from approximately 25 km to 55 km), but no other variables significantly affected fidelity. Our findings suggest that western Hudson Bay polar bears show a moderate amount of fidelity to denning areas, but not necessarily to specific sites.

## 1 Introduction

Site fidelity (i.e., the tendency to revisit previously occupied spatial locations; Switzer, 1993) has important consequences for animal population structure, distributions, and resilience to environmental change (Piper, 2011; Merkle *et al*., 2015; Gerber *et al*., 2019). Site fidelity can be advantageous for foraging (Wirsing *et al*., 2018), breeding (Greenwood, 1980; Hoover, 2003), efficient movement (Stamps, 1995), and overall fitness and survival (Bradshaw *et al*., 2004; Bose *et al*., 2017), and can develop with experience, age, and memory (Pyle *et al*., 2001; Zubiria Perez *et al*., 2021). However, the benefits of spatial fidelity are largely dependent on the spatiotemporal variability and predictability of the environment. In habitats with high predictability, site fidelity can be advantageous, as individuals can return to known high-quality habitat (Switzer, 1993; Morrison *et al*., 2021), and limit energy expenditure to explore potentially unprofitable new habitat (Gerber *et al*., 2019). Conversely, when habitats are unpredictable, the benefits of site fidelity are diminished and it can even be maladaptive if it prevents animals from optimally exploiting their environment (Switzer, 1993) or responding to environmental changes (Merkle *et al*., 2015). Therefore, patterns of fidelity have far-reaching implications for animal condition and survival (Bradshaw *et al*., 2004; Morrison *et al*., 2021).

Migratory animals use advanced spatial memory and exogenous cues to navigate their environment and take advantage of seasonal peaks in resource availability (Avgar *et al*., 2014). The evolution of seasonal migration has been hypothesized as primarily driven by maintenance of site fidelity to advantageous breeding locations (Winger *et al*., 2019). Such behaviour is notable in Arctic marine mammals, who display high seasonal site fidelity, migrating between habitats based on sea ice conditions (Tynan & DeMaster, 1997; Hauser *et al*., 2017; Shuert *et al*., 2022). Females can show higher spatial fidelity than males (e.g., Lone *et al*., 2013), potentially to garner energetic and reproductive benefits for gestation or care of young (Gauthier, 1990). Therefore, female-bias in site fidelity is more likely to be centred on reproductively important areas, such as parturition sites (Kelly *et al*., 2010). Arctic sea ice is highly variable, with the extent and configuration changing over days, months, and between years in response to seasons, weather, and climate change (Gough *et al*., 2004; Barber & Hanesiak, 2004; Comiso, 2006; Cavalieri & Parkinson, 2012). Migratory behaviour in Arctic marine mammals is therefore a balance between site fidelity to areas that fulfil important life history activities and optimizing energetic balance.

Polar bears (*Ursus maritimus*) are a migratory ice-obligate species that use sea ice for foraging, migration, and mating (Derocher *et al*., 2010; Stirling *et al*., 2016; Togunov *et al*., 2017; Bohart *et al*., 2021), but primarily use terrestrial maternal den sites in snow and peat banks to give birth and rear cubs in the first months of life (Harington, 1968; Florko *et al*., 2020). Polar bears mate on the sea ice in spring (Stirling *et al*., 2016), and implantation is delayed until autumn with parturition in early winter and den emergence in spring when females with newborn cubs return to the sea ice (Derocher *et al*., 1992). Denning areas have been recognized as critical habitat (Florko *et al*., 2020) and targeted for protection under the 1973 Agreement on the Conservation of Polar Bears (Prestrud & Stirling, 1994). Across all age classes, male and female polar bears exhibit general spatial fidelity (Derocher & Stirling, 1990; Mauritzen *et al*., 2001; Cherry *et al*., 2013; Lone *et al*., 2013), and females show fidelity to denning areas (Ramsay & Stirling, 1990; Amstrup & Gardner, 1994; Zeyl *et al*., 2010).

Polar bears have been shown to have some plasticity in denning locations, and this may have important implications in the context of climate change (Zeyl *et al*., 2010; Escajeda *et al*., 2018). As sea ice freeze-up and break-up patterns change through time (Sahanatien & Derocher, 2012; Castro de la Guardia *et al*., 2013; Kowal *et al*., 2017), it is possible that high site fidelity could have energetic consequences if bears do not adapt denning locations to correspond with new sea ice patterns (Derocher *et al*., 2004). Therefore, site fidelity in polar bears was predicted to decline with climate change (Cherry *et al*., 2013). Pregnant female polar bears fast while on land in WH for up to 8 months but come ashore with fat reserves that are used for body maintenance, gestation, and early lactation (Watts & Hansen, 1987; Ramsay & Stirling, 1988; Derocher & Stirling, 1998). We hypothesize that den site fidelity may be advantageous for pregnant females given their high body mass and consequent high energetic costs of movement to locate new and suitable denning habitat.

In this paper, we examine maternal den site fidelity of polar bears in the Western Hudson Bay sub-population (WH) in Manitoba, Canada. Hudson Bay is a seasonal sea ice system, in which all bears come ashore during the ice-free period (Derocher & Stirling, 1990; Cherry *et al*., 2013). Female polar bears have been using the same broad denning area for centuries (Hearne, 1795; Scott & Stirling, 2002), and there is evidence that they show general fidelity to a denning area but not necessarily the same den sites (Ramsay & Stirling, 1990). Some individual den sites have been reused for over 29 years but it is unknown if by the same individuals (Scott & Stirling, 2002). Den site fidelity in WH has not been quantified over a period of rapid environmental change, which may increase plasticity in denning behaviour. Sea ice across the Arctic is undergoing significant changes, and is generally declining in all 19 polar bear subpopulations across the Arctic (Stern & Laidre, 2016). In the WH subpopulation, polar bears have been shown to alter the their on-shore migrations with spatiotemporal changes in sea ice break-up (Cherry *et al*., 2013), which may subsequently influence chosen den locations. The objectives of our study are to: (i) quantify the degree of den site fidelity among denning females using capture and telemetry data, (ii) examine temporal, demographic, and environmental (i.e., spatial and temporal patterns of sea ice break up) factors that affect fidelity, and (iii) understand general directional patterns of inter-den movements.

## 2 Methods

### 2.1 Study area and population

WH polar bears use the sea ice of the Hudson Bay, which starts to form in November and melts entirely by August (Saucier *et al*., 2004; Klappstein *et al*., 2020). The majority of the population spends the ice-free period in Manitoba south of Churchill and the core of the denning area occurs in Wapusk National Park (Fig. 1). Unlike other subpopulations where females den close to shore (Florko *et al*., 2020), WH females will often move inland up to 80 km to find maternal den sites, possibly to avoid males that congregate along the shoreline (Jonkel *et al*., 1972; Derocher & Stirling, 1990; Richardson *et al*., 2005). They use maternity dens primarily in peat banks with high vegetation for den stability along riparian areas (Jonkel *et al*., 1972; Richardson *et al*., 2005). When snow accumulates sufficiently, females usually dig dens in the snow near the peat den (Ramsay & Stirling, 1990).

**Figure 1:**
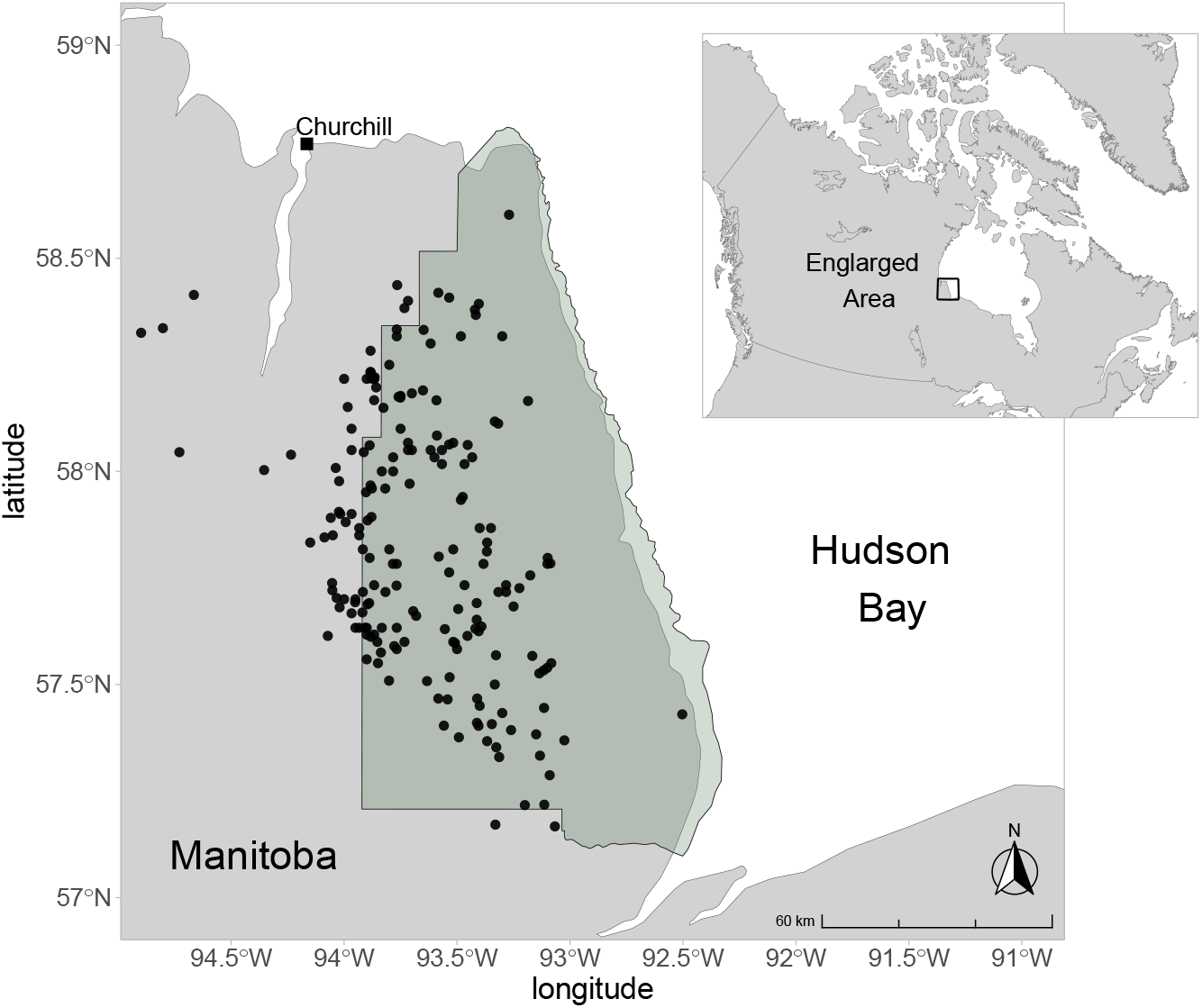
Adult female polar bear capture locations (black dots) in the western Hudson Bay denning area. Green shaded area is Wapusk National Park.

### 2.2 Den site data

We used capture-recapture data, as well as geographical positioning system (GPS) telemetry data to identify den sites. Bears were located from a helicopter and immobilized, following standard protocols (Stirling *et al*., 1989). Capture-recapture locations from 1979 *−* 2018 were obtained for lone adult female polar bears (*≥* 5 years old) in autumn (August – November) and adult females with 3 *−* 4 month old cubs-of-the-year captured in spring (late February – March) leaving the denning area. Captures were not performed specifically to assess denning, therefore we only retained captures as den site locations when they met at least some of the following criteria (in order of importance): i) autumn and spring locations of the same denning event matched, ii) a bear was found to be inside or at a den, and iii) a bear was recorded to be near a den (see Table A1 of Appendix A for a full description of den identification).

We also identified dens from GPS data from 2004 *−* 2019. Adult female polar bears were fitted with GPS collars programmed to provide locations every 4 hours, which either released automatically after 2 years or were removed upon recapture. To identify denning bears, we isolated locations that were on land between January and March, as this likely indicated denning. We only retained bears with *≥* 10 locations that we could use to identify a den through visual examination of their locations (Figure A1). To assess site fidelity, we only kept GPS-identified dens from individuals with another identified den, and locations within the core capture area (defined as the 100% minimum convex polygon of all capture locations). This removed two dens outside of the core denning area (Figure A1).

### 2.3 Quantifying den site fidelity

We calculated inter-year distance (denoted by *D*) as the Euclidean distance between two den sites as a metric of fidelity (Morrison *et al*., 2021). For all dens, we defined the year *t* based on the denning event, where a location in the autumn of a given year is considered to the be the same year as a location in spring of the following year (as they correspond to the same denning event). Then, for a den site ***x*** (bivariate location with easting/northing) in year *t*, the inter-year distance is calculated to the next capture,

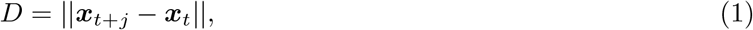

where *j* is the number of years between captures. We calculated all within-individual distances between successive locations for each individual (denote the vector as ***D***_*w*_). To statistically assess whether bears exhibit site fidelity, we compared ***D***_*w*_ to between-individual distances (denoted ***D***_*b*_). The vector ***D***_*b*_ was calculated as all unique pairs of locations of different individuals within successive years only. We considered ***D***_*b*_ to be a proxy for what would be expected at random within the current sampling design (i.e., if there were no site fidelity). We used non-parametric bootstrapping (i.e., sampling with replacement) to assess how ***D***_*w*_ differed from ***D***_*b*_. For each bootstrap iteration *i ∈ {*1, 2, …, *B}*, we sampled a new 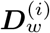 and 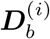, from which derived the sample means ( 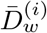 and 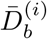) and difference in means 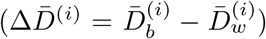. We used *B* = 10, 000 bootstrap iterations. We derived 95% confidence intervals as the 0.025 and 0.975 percentile of the bootstrapped means (and difference in means, where a 95% CI that does not overlap zero indicates a statistically significant difference between bootstrapped samples).

### 2.4 Factors affecting den site fidelity

We used a generalized additive model (GAM) to assess what factors influence the degree of site fidelity. GAMs are a special case of generalised linear models, in which the linear predictor can include smooth functions of covariates (most commonly smoothing splines; Wood, 2017; Pedersen *et al*., 2019). These smoothing splines are composed as the sum of *K* smaller basis functions, and a smoothing parameter *λ* controls the complexity (i.e., wiggliness) of the spline (Wood, 2017). Ultimately, *λ* controls the trade-off between constructing a smooth function and fitting the data (where *λ → ∞* indicates a straight line; Wood, 2017). Although some bears were captured multiple times, we did not consider a mixed effects model because 68% of bears only had a single *D*_*w*_ (and less than 8% had more than 4; see Results for details). Therefore, a mixed model may have been overly complex for our moderate sample size.

We considered the GAM response to be the within-individual *D*_*w*_, such that for observation *i*,

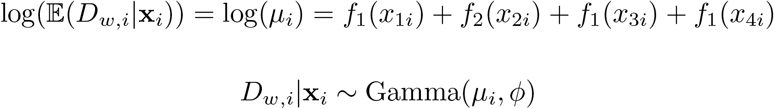

where *ϕ* is a scale parameter (related to the variance of the gamma distribution), *x*_1_ is years between captures, *x*_2_ is bear age, *x*_3_ is the difference in break up timing between successive captures (measured in days), and *x*_4_ is the difference in remnant ice location (measured in km). We considered the mean age between den locations, as fidelity is likely the result of the experience of the bear at both locations. Break up timing was defined as the day of the year when sea ice (with ice concentration *≥* 30%) was *≤* 10% of the maximum ice extent in winter across Hudson Bay. Remnant ice location was the centroid of the largest patch of ice on this break up date (McGeachy *et al*., in review). Based on the Pearson correlation coefficient *ρ*, no covariates were highly correlated (|*ρ*| was always less than 0.19; Figure B1, Appendix B). We set *K* = 5 (i.e., the number of basis functions) for all covariates. To conduct variable selection, we modelled all covariates with thin plate regression splines with shrinkage. These splines use specialised basis functions, where *λ* not only penalises towards smoothness but can also shrink functions toward zero, allowing covariates to be effectively removed if they do not improve model fit (Marra & Wood, 2011). Note that this approach will first penalise the function to a straight line, and then additional penalisation shrinks the function to zero (i.e., removing the variable). We estimated the model parameters with restricted maximum likelihood estimation using the `R` package `mgcv` (Wood, 2017).

### 2.5 Directionality of inter-den movements

In addition to distance, we wanted to better understand general directionality of bears’ inter-den movements, so we calculated the bearings (relative to north) of all successive den locations (denoted *ψ*_*d*_). Visual inspection of *ψ*_*d*_ suggested that there were two main directions. Therefore, we fitted a two-component von Mises mixture distribution (i.e., that allows for two mean directions) using the `movMF` package (Hornik & Grun, 2014). Then, we assessed whether the directionality between dens was influenced by the last remnant ice location. We calculated the bearings of the remnant ice locations between years (*ψ*_*r*_; relative to north), and assessed for correlation between *ψ*_*r*_ and *ψ*_*d*_ with a circular version of the Pearson’s product moment correlation. This test assesses for correlation between two circular variables and was implemented with the `circular` package in `R`. We recognized that inter-den movements may be related to inter-year ice movements, beyond a simple correlation. Therefore, we also calculated the angle of the successive den locations relative to the angle of the ice (denoted *ψ*_*d,r*_) and fitted a three-component von Mises mixture distribution to *ψ*_*d,r*_.

## 3 Results

We analysed 188 den locations from 78 individuals (*n* = 53 with 2 dens, *n* = 19 with 3 dens, *n* = 5 with 4 dens, and *n* = 1 with 5 dens). Of these, 111 were autumn captures, 63 were spring captures, and 14 were identified from GPS telemetry. Denning females had a median age of 16 years (range: 5 *−* 28 years), and the median year of denning was 1993 (range: 1980 *−* 2019).

### 3.1 Quantifying den site fidelity

We calculated all empirical inter-year distances, which resulted in a total of *n* = 111 within-individual *D*_*w*_ measurements that ranged from 1.7 *−* 107 km with mean *±* SD of 31.0 *±* 23.9 km. Empirical between-individual distances *D*_*b*_ were calculated between all possible successive inter-individual locations (*n* = 1242), and ranged from 0.3 *−* 158 km with a mean *±* SD of 49.5 *±* 28.8 km (18.5 km larger than the mean *D*_*w*_; Figure 2a). We assessed whether this difference between *D*_*b*_ and *D*_*w*_ was significant via non-parametric bootstrapping with 10,000 iterations. We estimated the bootstrapped metrics (with 95% CIs) as 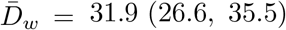, 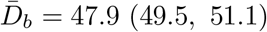 km (Figure 2b), and 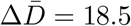 (13.8, 23.2) km (i.e., the 95% Cis did not overlap zero; Figure 2c).

**Figure 2:**
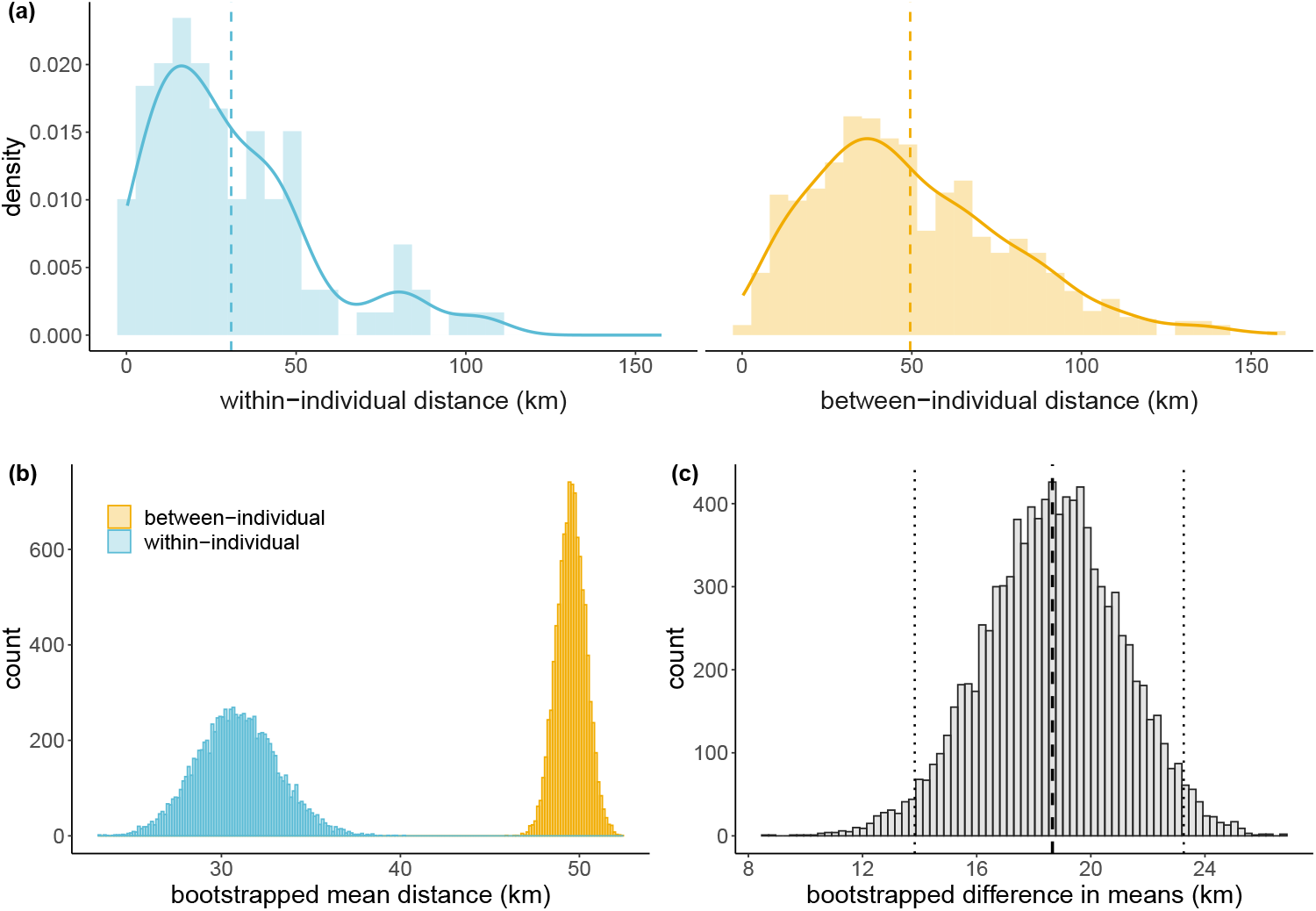
Comparison of within-individual and between-individual inter-year distances (D; km). (a) Empirical histograms with overlaid densities (estimated with Gaussian kernel density estimation, bandwidth of 8) and empirical means (dashed lines). The bottom panels show histograms of the bootstrapped metrics from 10,000 samples: (b) Mean within-individual 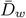 and mean between-individual 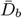 (both in km); and (c) Bootstrapped difference in means calculated as 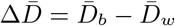 (dashed line is the mean and dotted lines are 95% CIs).

### 3.2 Factors affecting fidelity

For all GAM terms, higher effective degrees of freedom (EDF) indicate a more complex function (EDF = 1 indicates a straight line and EDF = 0 indicates removal from the model). All terms had EDF *<* 1 indicating that each term had a smoothing parameter *λ* large enough to shrink the effect towards zero, but the degree of shrinkage varied between covariates (Table 1). Years between captures had EDF close to 1 and results indicated a significant linear effect, whereas the other covariates were effectively shrunk to zero effect (Table 1, Figure 3). Additionally, the EDF for each term was well below *K* = 5, indicating that we used enough basis functions to allow sufficient complexity (i.e., wiggliness). Model diagnostics did not indicate any major concerns with fit (Appendix B), and our model explained 6.3% of the deviance.

**Table 1:** Results from a generalised additive model with a gamma distribution and a log link function. All covariate effects were estimated with thin plate regression splines with shrinkage, allowing effects to be shrunk to zero. K is the basis dimension (set prior to model fitting), EDF is the estimated effective degrees of freedom, and bolded p-values indicate statistical significance (α = 0.05).

| Covariate | $K$ | EDF | $p$-value |
| --- | --- | --- | --- |
| Years between captures | 5 | 0.88 | **0.006** |
| Mean age | 5 | 0.0002 | 0.46 |
| Difference in remnant ice location (km) | 5 | 0.02 | 0.30 |
| Difference in breakup dates (days) | 5 | 0.0004 | 0.63 |

**Figure 3:**
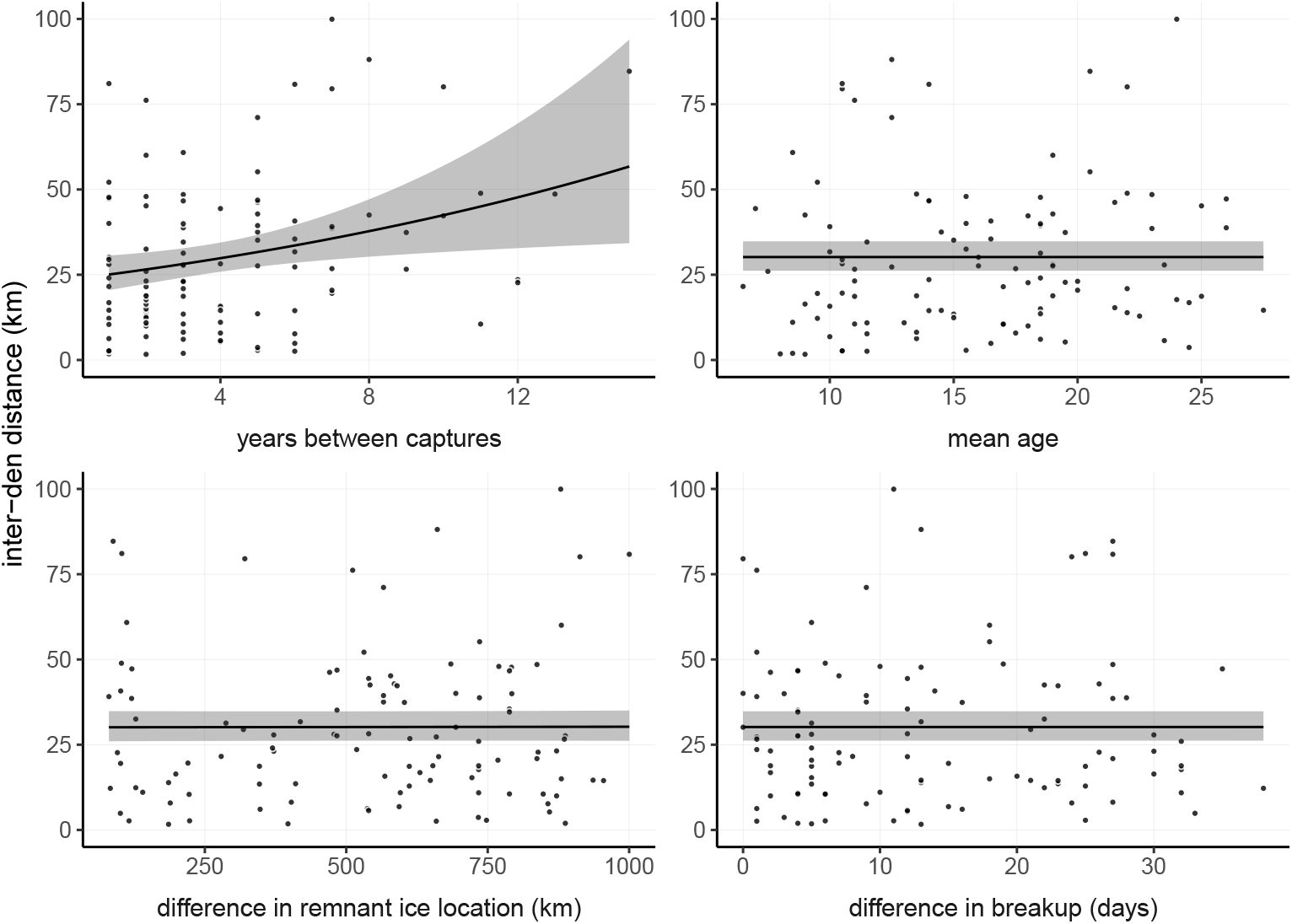
Predicted relationship between all model covariates and inter-year den distances, from a generalised additive model with a gamma distribution and a log link function. All effects were estimated with thin plate regression splines with shrinkage, allowing effects to be shrunk to zero. Black lines are the predicted values (with all other covariates held to their mean values), shaded bands are 95% across-the-function confidence intervals, and dots are the observed data.

### 3.3 Directionality of inter-den movements

A mixture von Mises distributions with *p* components is parametrised by its component means ***µ*** = *{µ*_1_, *µ*_2_, … *µ*_*p*_*}* and angular concentrations ***κ*** = *{κ*_1_, *κ*_2_, … *κ*_*p*_*}* (i.e., a measure of variation). We fitted a model with *p* = 2 to inter-den bearings *ψ*_*d*_, and found 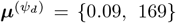 and 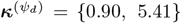 Figure 4a). This indicates that bears moved bi-modally between dens (generally north or south), but the southward movements had more angular concentration (i.e., they were more directional).

**Figure 4:**
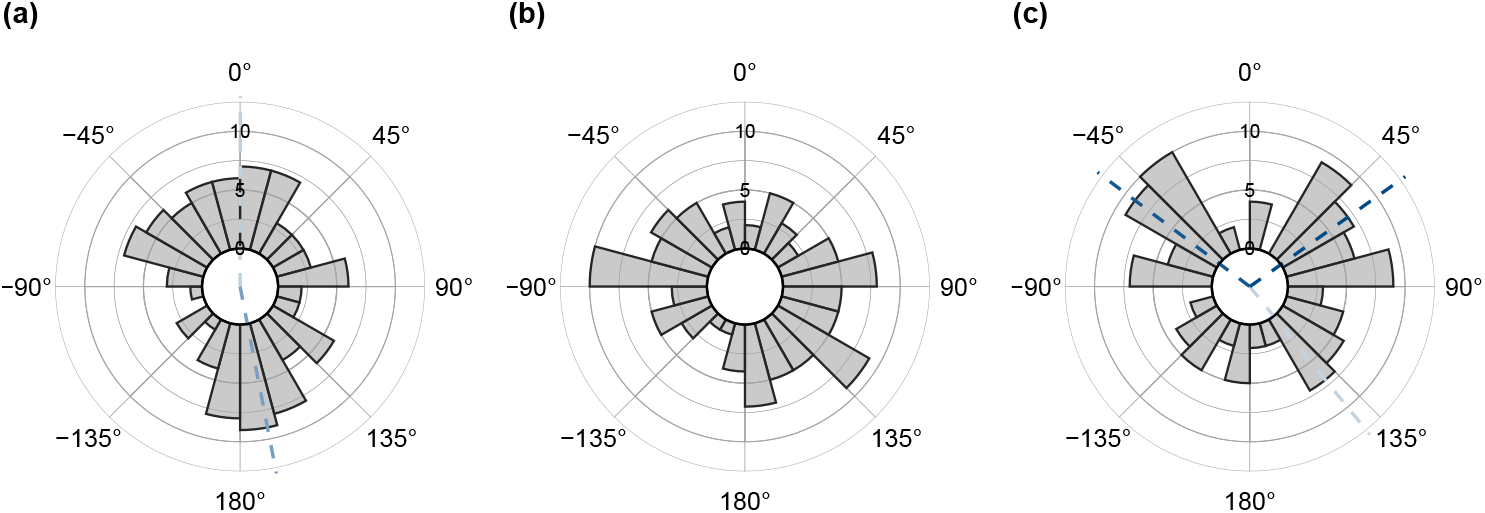
Histograms of bearings: (a) bearings of inter-den movements (where north is 0 degrees, (b) bearings between last remnant ice locations (where north is 0 degrees), and (c) angles of inter-den movements relative to inter-year remnant ice movements (where 0 indicates that inter-den movements are in the same direction as the ice). Dashed lines are the means of the von Mises mixture distributions, where darker lines have a higher angular concentration κ.

We assessed the correlation between *ψ*_*d*_ and inter-year ice bearings *ψ*_*r*_, and found no significant correlation (*ρ*(*ψ*_*d*_, *ψ*_*r*_) = *−*0.17, *p* = 0.07; Figure 4a,b). We complemented this analysis by fitting a three-component von Mises distribution to the angle of inter-den movement relative to that of remnant ice *ψ*_*d,r*_, which had 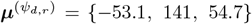 and 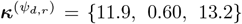. This indicates that inter-den movements tended to be neither directly with (0 degrees) or against (180 degrees) the ice direction, and angles at approximately 53-54 degrees in either direction are most concentrated (Figure 4c).

## 4 Discussion

Our analyses provided a 40 year assessment of maternal den site fidelity in western Hudson Bay. We quantified the general degree of fidelity, assessed factors affecting fidelity, and explored the directionality of inter-den movements. We found that female polar bears show a moderate amount of fidelity to den sites, based on a comparison of within-individual to between-individual inter-den distances. The level of fidelity was affected by time between denning events, but maternal age and sea ice conditions had no effect on the distance or direction between den sites. However, our findings are predicated on pregnant females returning to the sampling area as they would not be sampled if they denned in another area. Although this limits our inferences about fidelity to the core denning area, it does not preclude examination of within-area fidelity. Long-term satellite telemetry tracking in WH show that fidelity of females to on land areas is high (Stirling *et al*., 2004; Cherry *et al*., 2013) and the satellite tracked bears examined in our study showed fidelity to the study area. Further, we largely relied on capture data, and assumed that bears captured in autumn were pregnant and at their final denning site, and that bears caught in spring had not already moved from their den before capture. Although there was some uncertainty in the capture data, we attempted to mitigate this issue by only retaining captures with the most reliable denning information (i.e., locations of bears that were close to maternal dens and/or had matching autumn and spring locations of the same denning event/year). If a small subset of the data were misclassified, they still contained information about on-land site location and were still useful in assessing general spatial fidelity.

We found that female polar bears denning in WH showed a moderate amount of site fidelity. Within-individual distances were approximately 18.5 km shorter than between-individual distances (and this difference was found to be significant via non-parametric bootstrapping). This is a potentially meaningful difference on the scale of bootstrapped inter-den distances 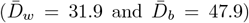. This observed fidelity may be due to philopatry, on-ice spatial fidelity, and a tendency to travel a similar distance in-land once returning to shore. Spatial fidelity of female ursids to the general area of their birth site is in part associated with higher levels of philopatry in females than in males (Manchi & Swenson, 2005; Roy *et al*., 2012; Takayama *et al*., 2023). Similarly, philopatry in polar bears appears common across both sexes (Derocher & Stirling, 1990; Paetkau *et al*., 1999; Zeyl *et al*., 2010) but may be lower in males that move across subpopulations more often (McGeachy *et al*., in press). This philopatry may weaken with time, indicated by the result that *D*_*w*_ tended to (modestly) increase with time between captures. That is, *D*_*w*_ was predicted to increase from approximately 25 km (at 1 year between captures) to 55 km (at 15 years between captures), which may suggest that fidelity to previous sites sometimes weakened as bears shifted their denning locations through time.

We expected den fidelity to be driven by the similarity in sea ice conditions between denning years, but we found no effect of sea ice conditions on inter-den distance or directionality. Directionality between dens used by individuals were somewhat oriented to the north and southeast, which is broadly parallel to the Hudson Bay coastline. Previous research found that WH maternity dens range from 12.6 to 80.0 km from the coast (Richardson *et al*., 2005). Our findings are consistent with bears selecting dens approximately the same distance from the coast, but moving northward or southward between denning years. Sea ice break up patterns in Hudson Bay affect the distribution of polar bears (Stirling *et al*., 2004; Cherry *et al*., 2013) so we expected that the observed directional shifts may be related to on land arrival locations of pregnant females. Although we found no effect of sea ice conditions on the distance or directionality between den sites, these sea ice covariates were only approximations of where and when bears may come ashore. We lacked information on the exact location of on-land arrival, which may be more related to bears’ chosen den sites.

Although we observed some evidence of fidelity, when considered in the context of polar bears’ large home ranges, this may be considered a modest difference. Polar bears have home ranges much larger than expected for mammals of their size (Ferguson *et al*., 1999) and adult females in WH have annual home ranges > 260, 000 km^2^ (McCall *et al*., 2015). Similar to other polar bear subpopulations (Andersen *et al*., 2012; Florko *et al*., 2020), suitable den habitat is widely available in the core denning area in WH (Clark *et al*.,1997; Richardson *et al*., 2005). This may partially explain why *D*_*w*_ did not increase with mean age, which may be expected if bears learn about the spatial distribution of suitable denning habitat (Morrison *et al*., 2021); this learning behaviour may not be necessary in WH, where suitable habitat is widely distributed.

Further, movements to return to previous dens may yield little benefit given the abundance of denning habitat, particularly considering polar bears’ high locomotory costs. Early energetics studies found that polar bears were inefficient in movement with the costs more than twice that predicted for a quadruped of their size (Øritsland *et al*., 1976; Hurst *et al*., 1982), but more recent research using biologgers found that costs were even higher than previously assumed (Pagano *et al*., 2018). Thus, the scale of den site fidelity is relatively small in the overall context of the space use of polar bears and energetic costs or benefits. Similarly, female polar bears with newborn cubs departing the WH den area in spring show strong orientation to the northeast towards foraging areas and noted that the bears did not take the shortest path to the coast, indicating that not all movements are driven by energetics (Ramsay & Andriashek, 1986; Yee *et al*., 2017). Inter-individual variability in den fidelity may be related to observed inter-individual variability in energetic strategies (observed in the Beaufort Sea subpopulation; Klappstein *et al*., 2022). On the scale of annual home range size, the differences in migration distance to the coast or foraging areas may be minimal.

Broadly, polar bear maternity den fidelity appears similar to congenerics with fidelity to general denning areas, but not specific locations, as documented in brown bears (*U. arctos*) (Manchi & Swenson, 2005; Sorum *et al*., 2019) and Asian black bears (*U. thibetanus*) (Kozakai *et al*., 2017). Within the context of polar bear movements, energetics, space use, and site fidelity, the costs and benefits of den site fidelity remain unclear. Longer-term monitoring of den site use based on telemetry data may allow insights into environmental factors that influence site selection (e.g., sea ice break-up patterns, on land arrival locations). Additionally, further research with more detailed individual-specific biological factors, such as age, experience, reproductive success, and body condition may yield new insights on site fidelity for maternity dens. Lastly, we believe our analytical approach is a useful means of assessing site fidelity and has application to other species where location data is available.

## 5 Acknowledgements

This project was supported by the Banrock Station Environmental Trust, Canadian Association of Zoos and Aquariums, the Churchill Northern Studies Centre, Canadian Wildlife Federation, Care for the Wild International, Earth Rangers Foundation, Environment and Climate Change Canada, Hauser Bears, the Isdell Family Foundation, Kansas City Zoo, Manitoba Department of Agriculture and Resource Development, Manitoba Sustainable Development, Natural Sciences and Engineering Research Council of Canada, Parks Canada Agency, Pittsburgh Zoo Conservation Fund, Polar Bears International, Polar Continental Shelf Project, Quark Expeditions, San Diego Zoo Wildlife Alliance, Schad Foundation, the University of Alberta, World Wildlife Fund Canada, and Wildlife Media Inc.

## 6 Ethics statement

Capture and handling protocols were approved by the University of Alberta BioSciences Animal Policy and Welfare Committee and Environment and Climate Change Canada, Prairie and Northern Region Animal Care Committee in accordance with the Canadian Council on Animal Care guidelines. Research was permitted by the Government of Manitoba and the Parks Canada Agency.

## A Additional details of den data

**Table A1:** Possible denning locations were manually classified based on our confidence that a bear was at its denning location. This was based on several pieces of information, including whether the bear was located at a den, movement between successive captures in a single season, movement between seasons, etc. N refers to the number of dens in our final dataset with each confidence level; we only included bears with confidence levels ≥ 4.

| confidence level | criteria | $N$ |
| --- | --- | --- |
| 1 (lowest) | Travelling/dead bears | - |
| 2 (very low) | Travelling bears, little information | - |
| 3 (low) | Lacking information | - |
| 4 (moderate) | Indication of den, with ambiguity (single location; discrepancy between locations) | 35 |
| 5 (moderate-high) | Likely at a den, but some ambiguity in the records or only single location | 105 |
| 6 (high) | Similar to level 5, but with more reliable information or multiple locations | 13 |
| 7 (highest) | Almost certainly at den (GPS; agreement between multiple locations) | 35 |

**Figure A1:**
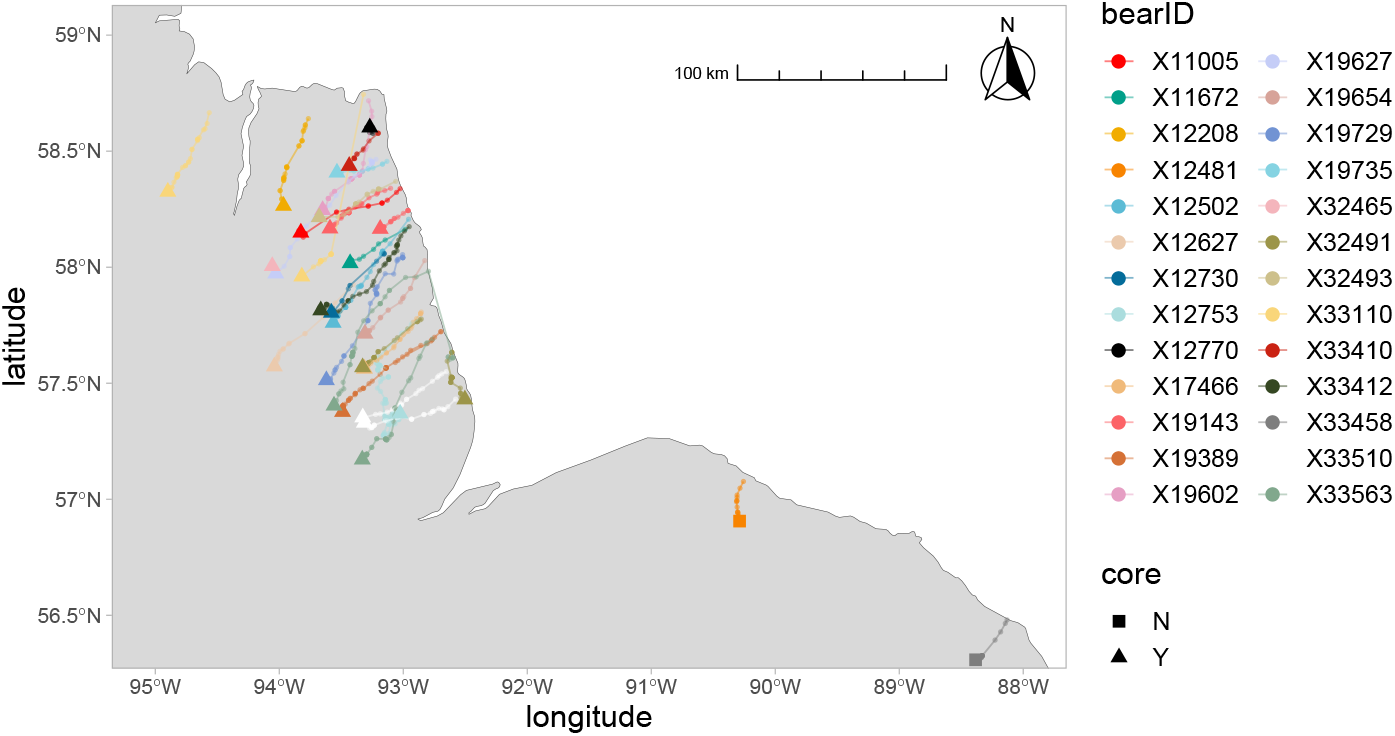
GPS tracks of denning bears. “Core” refers to whether or not the den location was within the main/core denning area and therefore included in the analysis.

## B Additional details of statistical analyses

**Figure B1:**
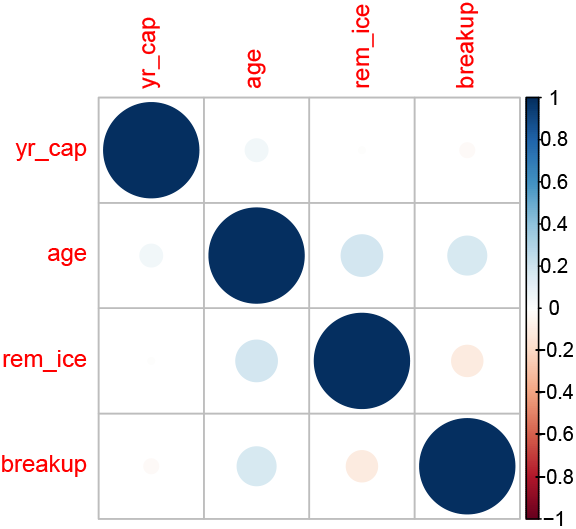
Correlation between covariates included in the generalised additive model described in the main text: years between captures (yr_cap), mean age (age), difference in remnant ice location (rem_ice), and difference in breakup date (breakup). The circle colour/size corresponds to the direction/size of the Pearson’s correlation coefficient.

**Figure B2:**
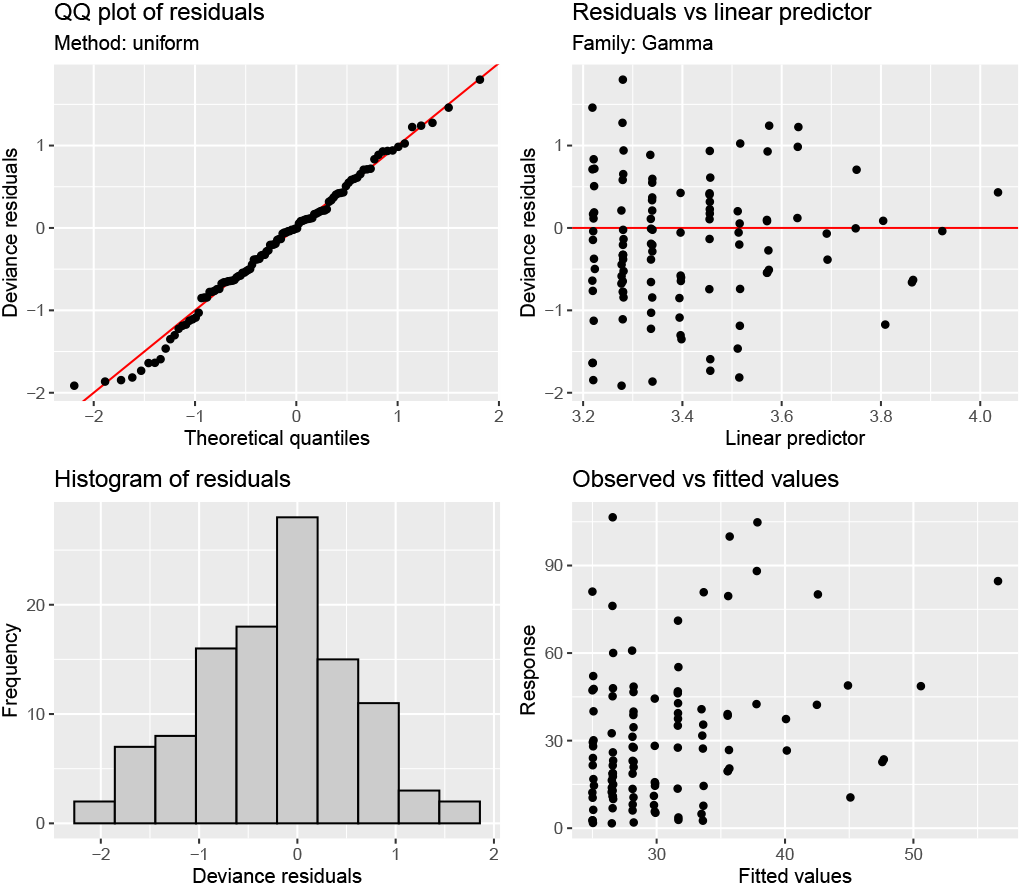
Diagnostic plots from the generalised additive model described in the main-text, produced via the `appraise` function of the package `gratia`.

